# Adult *Camk2a* gene reinstatement restores the learning and plasticity deficits of *Camk2a* knockout mice

**DOI:** 10.1101/2022.06.14.496142

**Authors:** Pomme M.F. Rigter, Ilse Wallaard, Mehrnoush Aghadavoud Jolfaei, Jenina Kingma, Laura Post, Minetta Elgersma, Ype Elgersma, Geeske M. van Woerden

**Author notes:** Address correspondence to: Ype Elgersma, Department of Clinical Genetics; Or to Geeske van Woerden, Department of Clinical Genetics and Department of Neuroscience, Erasmus Medical Center, Rotterdam, Dr. Molewaterplein 40, 3015 GD, The Netherlands.

## Abstract

With the recent findings that mutations in the gene encoding the α-subunit of calcium/calmodulin-dependent protein kinase II (CAMK2A) causes neurodevelopmental disorder (NDD), it is of great therapeutic relevance to know if there a critical developmental time window in which CAMK2A needs to be expressed for normal brain development, or whether expression of the protein at later stages is still beneficial to restore normal functioning. To answer this question, we generated an inducible *Camk2a* mouse model, which allows us to express CAMK2A at any desired time. Here, we show that adult expression of CAMK2A rescues the behavioural and electrophysiological phenotypes seen in the *Camk2a* knock-out mice, including spatial and conditional learning and synaptic plasticity. These results suggest that CAMK2A does not play a critical irreversible role in neurodevelopment, which is of importance for future therapies to treat CAMK2A-dependent disorders.

## Introduction

Intellectual disability (ID), a condition defined by an IQ below 70, is a predominant feature of neurodevelopmental disorders (NDDs) affecting approximately 1% of the global world population (1, 2). Due to next generation sequencing, many genes have now been implicated in this clinical condition (3, 4). One of the genes recently implicated in ID is *CAMK2A*, which encodes the alpha-subunit of calcium/calmodulin-dependent protein kinase II (CAMK2A), a highly abundant kinase in the brain that has been shown to play a critical role in hippocampus-dependent synaptic plasticity, learning and memory in mice (5, 6). In humans a similar role for CAMK2A has been suggested, as all individuals carrying mutations in *CAMK2A* suffer from a NDD with mild to severe ID, and some from Autism Spectrum Disorder (7–9).

With the diagnosis for the *CAMK2A*-dependent NDD established, doors open towards identifying a potential therapy. However, the important question that emerges is whether the genetic mutation causes alterations during early neurodevelopment which might be irrevocable, and thus dictate a critical window for therapeutic intervention (10, 11). Current literature shows that the answer to this question depends strongly on the gene involved in the NDD and the behaviour tested (12–18). For example in a mouse model for Rett syndrome, reinstatement of the gene *Mecp2* in adulthood improves all phenotypic behaviour, but seldom back to wild type performance (12, 13). However, in other models for NDDs the ability to rescue phenotypes was also shown to depend on the behaviour tested, as exemplified by *Grin1, Shank3, Ube3a* and *Syngap1* (14–18). Social and cognitive deficits, but not motor and anxiety deficits, are largely, but not completely rescued upon adult reinstatement of GRIN1 (14) and fully rescued upon SHANK3 reinstatement (15). In a mouse model for Angelman Syndrome, adult reinstatement of UBE3A does not rescue behavioural phenotypes found in the *Ube3a* mouse (16). However, reinstatement of UBE3A at postnatal day 21 (P21) rescues the motor skill deficits on the rotarod test, but not deficits in anxiety-related or intrinsic behaviour, suggesting an early critical window for these forms of behaviour (16). In a *Syngap1* haploinsufficiency model, cognitive behaviour cannot be rescued upon adult reinstatement, whereas motor and anxiety-related behaviour can only be rescued upon very young reinstatement, at P1 but not P21 (17, 18). Interestingly, pharmacological treatment with a GSK3-β inhibitor of the *Syngap1* mouse model during the critical period of P10-16 does rescue social, cognitive, and anxiety-related behaviour, and partially motor behaviour (19). Finally, the only phenotype that could be rescued upon adult reinstatement in all NDD mouse models discussed here is the electrophysiological plasticity (12, 15, 16, 20–22).

CAMK2A starts to be expressed around postnatal day 1, which means that it is not required for prenatal neurodevelopment (23). Whereas in the adult brain CAMK2A has been shown to play a critical role in normal brain functioning, as adult deletion of *Camk2a* is equally detrimental for learning and plasticity as germline deletion (24), not much is known about a possible critical period for CAMK2A during the early postnatal development and whether the phenotypes seen in adult mice could potentially be rescued upon CAMK2A expression.

To assess whether there is a role of CAMK2A in postnatal neurodevelopment and if a critical period for CAMK2A expression exists, we generated an inducible *Camk2a* mouse model, in which we can reinstate *Camk2a* at the time of our choosing. We show that adult reinstatement of *Camk2a* rescues all behavioural and electrophysiological phenotypes seen in the *Camk2a* knockout mice. These results indicate that absence of CAMK2A during development does not lead to irreversible alterations in brain development, and that potential therapies for CAMK2A-related disorders do not require specific early time windows.

## Results

### *Generation of an inducible* Camk2a *mouse*

To study the role of CAMK2A during neurodevelopment, we generated a novel mouse model. A transgenic cassette containing *Camk2a* exon 2 fused to the *tdTomato* gene with a transcriptional stop at the end, flanked with loxP sites, was inserted between exon 1 and 2 of the endogenous *Camk2a* gene (referred to as CAMK2 Lox – Stop – Lox (CAMK2^LSL^)), allowing for temporally controlled re-expression of *Camk2a* upon Cre-mediated deletion of the transgenic cassette. Heterozygous CAMK2^LSL^ mice were then crossed with the transgenic tamoxifen-inducible CAG-Cre^ESR^ line, to obtain both the CAMK2^LSL^ and the inducible CAMK2A^LSL^;CAG-Cre^ESR^ mouse line (Figure 1A). As control, wild type littermates negative or positive for Cre-expression were taken along (CAMK2A^WT^ and CAMK2A^WT^;CAG-Cre^ESR^, respectively).

**Figure 1:**
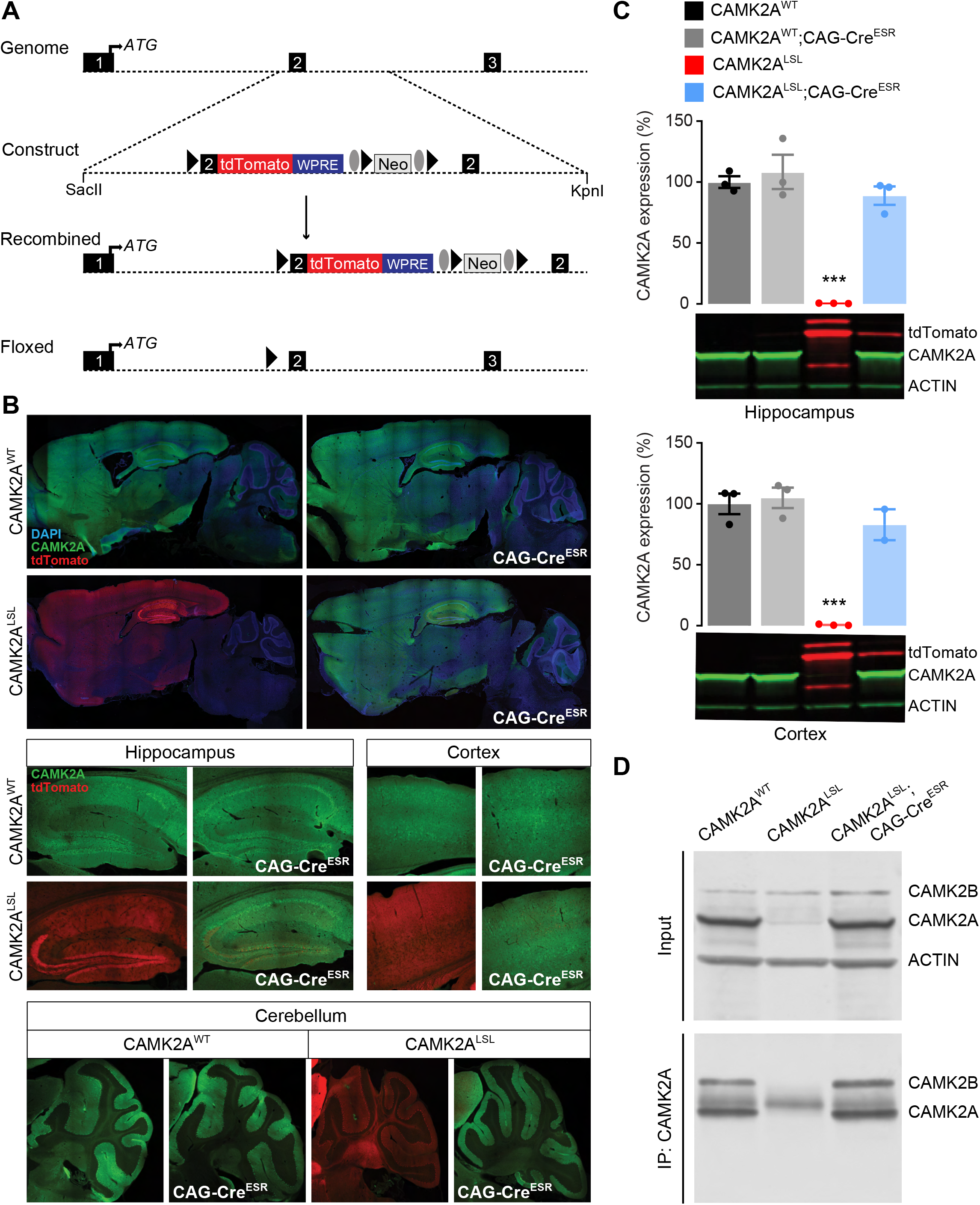
Generation and molecular analysis of the inducible CAMK2A^LSL^ ;CAG-Cre^ESR^ knock-in mouse. **A)** To generate CAMK2A^LSL^;CAG-Cre^ESR^ mice, a transgenic cassette was generated in which the *tdTomato* open reading frame was fused to exon2 of the *Camk2a* gene followed by the woodchuck hepatitis virus post-transcriptional regulatory element (WPRE) flanked by LoxP sites and a neomycin cassette, flanked by Frt sites, for positive selection. This cassette was inserted in the *Camk2a* gene in the genome through homologous recombination, as indicated in the schematic overview, which results in *tdTomato* expression under the control of the *Camk2a* promotor. Breeding with CAG-Cre^ESR^ line resulted in floxed Lox-P sites and transcription of the *Camk2a* gene. Triangles indicate LoxP sites, ovals indicate Frt sites. Restriction sites used for linearization of the plasmid are indicated. **B)** Representative images of immunofluorescent stainings for CAMK2A and tdTomato on sagittal slices of whole brain (top), hippocampus and cortex (middle), and cerebellum (bottom). **C)** Immunoblot analysis of hippocampal and cortical tissue probed with CAMK2A, tdTomato and Actin antibodies revealed significant CAMK2A reinstatement in the CAMK2A^LSL^;CAG-Cre^ESR^ mice (one-way ANOVA with Bonferroni’s post-hoc analysis. Hippocampus: F_(3,8)_ = 36.09, p < 0.0001, CAMK2A^WT^ vs CAMK2A^WT^;CAG-Cre^ESR^ p > 0.999, CAMK2A^WT^ vs CAMK2A^LSL^ p = 0.0002, CAMK2A^WT^ vs CAMK2A^LSL^;CAG-Cre^ESR^ p > 0.999, CAMK2A^LSL^ vs CAMK2A^LSL^;CAG-Cre^ESR^ p = 0.0004. Cortex: F_(3,7)_ = 41.85, p < 0.0001, CAMK2A^WT^ vs CAMK2A^WT^;CAG-Cre^ESR^ p > 0.999, CAMK2A^WT^ vs CAMK2A^LSL^ p = 0.0002, CAMK2A^WT^ vs CAMK2A^LSL^;CAG-Cre^ESR^ p > 0.999, CAMK2A^LSL^ vs CAMK2A^LSL^;CAG-Cre^ESR^ p = 0.0013). **D)** Immunoprecipitation of CAMK2A on cortical tissue of CAMK2A^WT^, CAMK2A^LSL^ and CAMK2A^LSL^;CAG-Cre^ESR^ mice. Representative blots probed with antibodies against CAMK2A, CAMK2B and Actin are shown. Band of IgG heavy chain is detectable in IP blot between the CAMK2A and CAMK2B bands. Data represents mean ± SEM; n = 2-3 per group; *** p < 0.001.

In the absence of Cre-driven recombination, CAMK2A expression in adult CAMK2A^LSL^ mice is abolished and replaced by tdTomato expression driven by the endogenous CAMK2 promotor (Figure 1B). Administration of tamoxifen (0.1 mg/ml, 4 consecutive days) to adult mice (>8 weeks old), successfully deleted the tdTomato transgene and induced CAMK2A expression throughout the brain (Figure 1B). To get insight in the level of CAMK2A protein expression upon reinstatement, a western blot was performed on hippocampal and cortical tissue of the mice 3-4 weeks after gene reinstatement. Western blot analysis revealed 89% and 83% of CAMK2A expression upon reinstatement in the hippocampus and cortex respectively of CAMK2A^LSL^;CAG-Cre^ESR^ mice, with low levels of tdTomato still present (Figure 1C; all statistics are given in the figure legends).

CAMK2A is known to form heteromeric holoenzyme complexes with CAMK2B consisting of 12-14 subunits (25, 26). Since we started to express CAMK2A here in the adult brain, having the mice to develop with only CAMK2B present, we tested whether newly expressed CAMK2A forms heteromeric holoenzymes again with CAMK2B. Four weeks after tamoxifen injections, we immunoprecipitated CAMK2A from cortex tissue and immunoblotted for both isoforms. CAMK2B was successfully pulled down in CAMK2A^WT^ and CAMK2A^LSL^;CAG-Cre^ESR^ mice, but not in CAMK2A^LSL^, indicating correct formation of heteromeric holoenzymes consisting of CAMK2A and CAMK2B subunits, upon CAMK2A reinstatement (Figure 1D). Note that the input showed similar CAMK2B expression in all groups, confirming previous findings (27) that loss of CAMK2A does not result in upregulation of CAMK2B.

### *Adult* Camk2a *reinstatement rescues learning and plasticity phenotypes*

*Camk2a* knockout mice are known to have severe impairments in spatial as well as associative learning (5, 27, 28). To confirm that our CAMK2A^LSL^ mice behave as a true knockout, and whether adult gene reinstatement can rescue these phenotypes, 12-16-week-old mice were injected with tamoxifen and 5 weeks later, spatial and associative learning was assessed using the Morris water maze and contextual and cued fear conditioning (Figure 2A).

**Figure 2:**
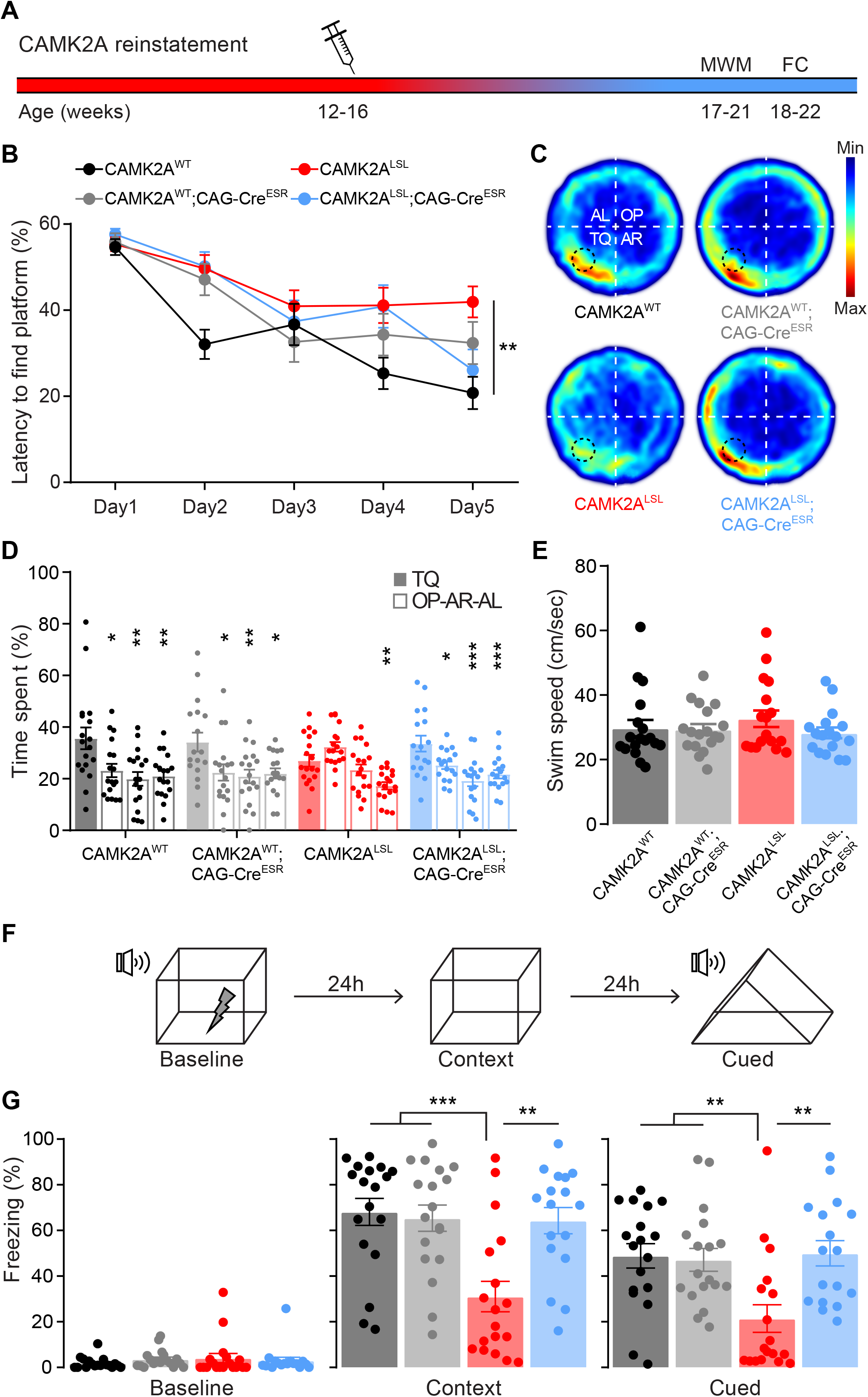
Adult CAMK2A expression rescues cognitive spatial and associative learning. **A)** Timeline representing reinstatement of CAMK2A in adulthood and behavioural testing, MWM = Morris water Maze, FC = Fear Conditioning. Injection needle indicates tamoxifen injections on 4 consecutive days. **B)** Latency to find the platform over 5 consecutive days, with two sessions per day (two-way repeated measures ANOVA with Bonferroni’s post-hoc analysis on genotype. Interaction time x genotype: F_(12,268)_ =2.44, p = 0.005; genotype: F _(3,67)_ =3.919, p = 0.0122, CAMK2A^WT^ vs CAMK2A^WT^;CAG-Cre^ESR^ p = 0.4251, CAMK2A^WT^ vs CAMK2A^LSL^ p = 0.0085, CAMK2A^WT^ vs CAMK2A^LSL^;CAG-Cre^ESR^ p = 0.1296, CAMK2A^WT^;CAG-Cre^ESR^ vs CAMK2A^LSL^ p = 0.8402, CAMK2A^LSL^ vs CAMK2A^LSL^;CAG-Cre^ESR^ p > 0.999). **C)** Heat map for probe trial on day 6 indicates where the mice spent most time in the Morris water maze for each genotype. For analysis the arena was divided into 4 quarters with the platform in the target quadrant (TQ). OP = opposite, AR = adjacent right, and AL = adjacent left quadrant. **D)** Time spent in each quadrant (TQ, OP, AR and AL, respectively) during probe trial on day 6, asterisks indicate differences compared to time spent in TQ within genotype (one-way ANOVA with Bonferroni’s post-hoc analysis. CAMK2A^WT^: F_(3,68)_ = 5.94, p = 0.0012, TQ vs OP p = 0.0128, TQ vs AR p = 0.0012, TQ vs AL p = 0.0027. CAMK2A^WT^;CAG-Cre^ESR^: F_(3,68)_ = 4.62, p = 0.0053, TQ vs OP p = 0.0166, TQ vs AR p = 0.0055, TQ vs AL p = 0.0127. CAMK2A^LSL^: F_(3,68)_ = 10.46, p < 0.0001, TQ vs OP p = 0.171, TQ vs AR p = 0.600, TQ vs AL p = 0.0024. CAMK2A^LSL^;CAG-Cre^ESR^: F_(3,64)_ = 8.20, p = 0.0001, TQ vs OP p = 0.0262, TQ vs AR p < 0.0001, TQ vs AL p = 0.0009). **E)** Swimming speed during probe trial (one-way ANOVA, F_(3,67)_ = 0.74, p = 0.530). **F)** Schematic overview of the fear conditioning procedure. **G)** Amount of time the mice spent freezing in baseline, context and cued condition (one-way ANOVA with Bonferroni’s post-hoc analysis. Baseline: F_(3,67)_ = 0.52, p = 0.671. Context: F_(3,67)_ = 8.47, p < 0.0001, CAMK2A^WT^ vs CAMK2A^WT^;CAG-Cre^ESR^ p > 0.999, CAMK2A^WT^ vs CAMK2A^LSL^ p = 0.0003, CAMK2A^WT^ vs CAMK2A^LSL^;CAG-Cre^ESR^ p > 0.999, CAMK2A^LSL^ vs CAMK2A^LSL^;CAG-Cre^ESR^ p = 0.0016. Cued; F_(3,67)_ = 6.24, p = 0.0008, CAMK2A^WT^ vs CAMK2A^WT^;CAG-Cre^ESR^ p > 0.999, CAMK2A^WT^ vs CAMK2A^LSL^ p = 0.0041, CAMK2A^WT^ vs CAMK2A^LSL^;CAG-Cre^ESR^ p > 0.999, CAMK2A^LSL^ vs CAMK2A^LSL^;CAG-Cre^ESR^ p = 0.0031. Data represents mean ± SEM, n = 17-18 mice per group. * p < 0.05, ** p < 0.01, *** p < 0.001.

Mice were trained to find the hidden platform of the Morris water maze for 5 days. Although latencies to find the platform decreased over time for all genotypes, CAMK2A^LSL^ consistently showed longer latencies compared to CAMK2A^WT^, suggesting impaired procedural learning (Figure 2C). As we removed the platform for the probe trial to assess spatial learning, CAMK2A^WT^ and CAMK2A^WT^;CAG-Cre^ESR^ mice showed clear preferences to spend time in the target quadrant, proving they successfully learned the location of the platform (Figure 2D). As expected, CAMK2A^LSL^ mice did not show any preference for the target quadrant confirming impaired spatial learning. Adult *Camk2a* gene reinstatement resulted in a complete rescue of the spatial learning deficits, as CAMK2A^LSL^;CAG-Cre^ESR^ mice showed clear preference for the target quadrant, equivalent to CAMK2A^WT^ mice (Figure 2D). Importantly, swim speed was similar between genotypes (Figure 2E).

For the fear conditioning paradigm, on day 1, mice were allowed to explore the fear conditioning box for 150 sec after which they were exposed for 20 sec to a tone, which ended simultaneously with a 2 sec mild foot shock. 24 hours later on day 2, mice were placed back in the same box and contextual memory was assessed by measuring the amount of time the mice showed freezing behaviour (Figure 2F). Confirming the previously published impaired associative learning (24, 28), CAMK2A^LSL^ mice spent significantly less time freezing compared to the other genotypes (Figure 2G). On day 3, mice were placed in a different context and were presented with the tone to test for amygdala-dependent cued learning. Again, CAMK2A^LSL^ mice showed impaired associative learning, spending significantly less time freezing compared to the other genotypes. Similarly as in the Morris water maze experiment, adult expression of CAMK2A induced a full rescue of the associative learning phenotype, as CAMK2A^LSL^;CAG-Cre^ESR^ mice showed equal freezing time compared to CAMK2A^WT^ mice both in contextual and cued fear conditioning (Figure 2G).

It is generally accepted that synaptic plasticity, i.e. the strengthening of synapses, underlies learning and memory (29). Indeed, long-term potentiation (LTP) at the Schaffer collateral-CA1 synapse in the hippocampus is known to be impaired in *Camk2a* knockout mice (6, 27, 30). With all the cognitive phenotypes being rescued upon adult *Camk2a* gene reinstatement, we hypothesized that also synaptic plasticity would be rescued upon adult reinstatement. We first measured basal synaptic transmission at the Schaffer collateral-CA1 synapse, where the CAMK2A^LSL^ showed similar strength compared to CAMK2A^WT^ (Figure 3, A-C). We did find a small Cre effect in the fEPSP of the input/output paradigm in the CAMK2A^WT^ mice (Figure 3B). Upon induction of LTP we found, consistent with previous findings (6, 27), a significant impairment in hippocampal LTP in the CAMK2A^LSL^ mice compared to the other genotypes. Additionally, confirming our hypothesis, LTP was completely normalised in the CAMK2A^LSL^;CAG-Cre^ESR^ mice, proving a rescue also of synaptic plasticity upon *Camk2a* gene reinstatement (Figure 3D and E).

**Figure 3:**
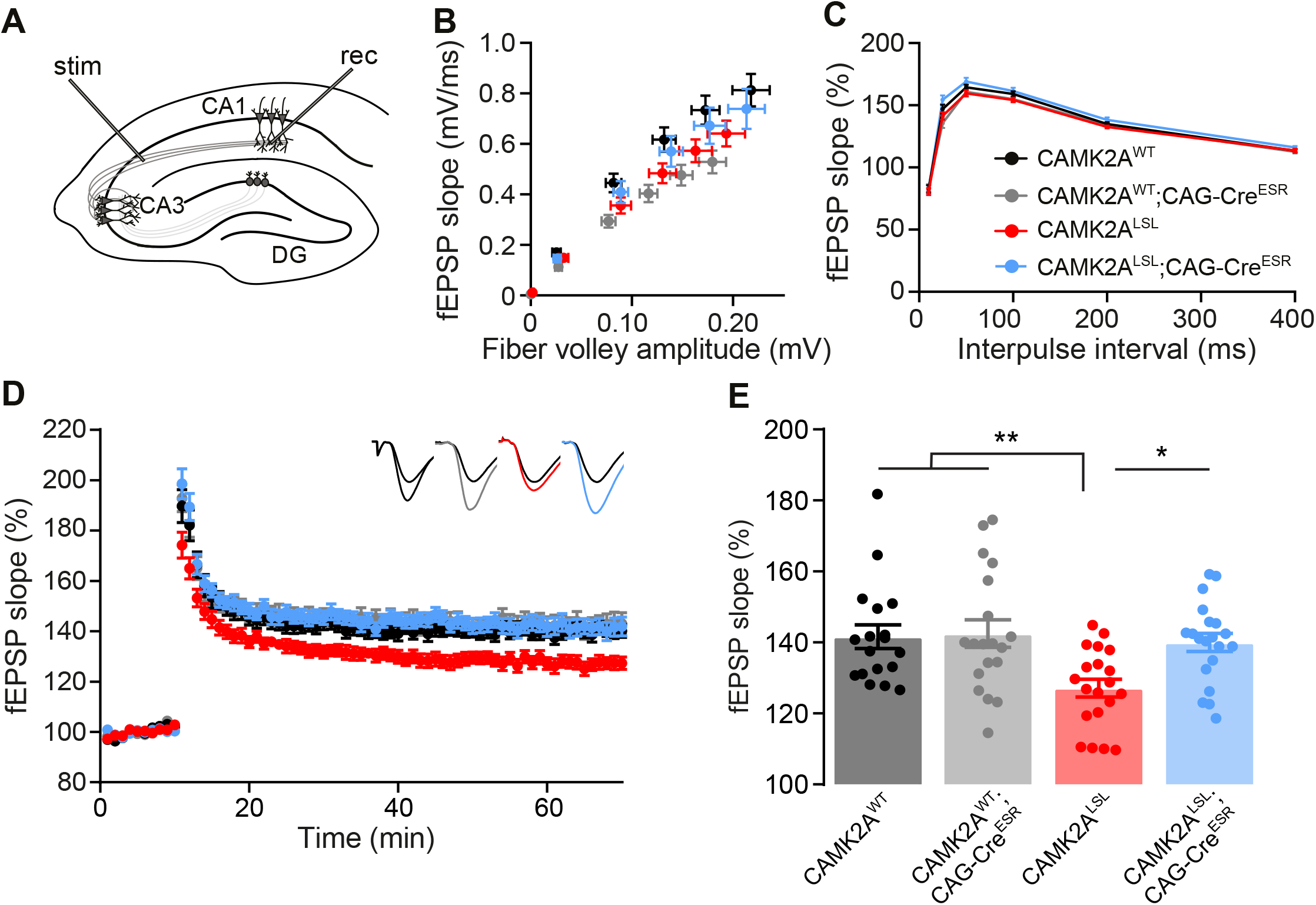
Adult CAMK2A expression rescues synaptic long-term plasticity. **A)** Schematic overview of the placement of the electrodes for the extracellular field recordings done in Schaffer collateral-CA1 synapses of hippocampal slices. **B)** Basal synaptic transmission, fiber volley and fEPSP slope recorded after increasing stimulation strength (10-100 mA; two-way repeated measures ANOVA with Bonferroni’s post-hoc analysis on genotype. Fiber volley: interaction time x genotype: F_(15,540)_ =1.40, p = 0.1401; genotype: F_(3,108)_ = 0.55, p = 0.6481. fEPSP: interaction time x genotype: F_(15,540)_ =4.18, p < 0.0001; genotype: F_(3,108)_ = 4.01, p = 0.0095, CAMK2A^WT^ vs CAMK2A^WT^;CAG-Cre^ESR^ p = 0.0001, CAMK2A^WT^ vs CAMK2A^LSL^ p = 0.0544, CAMK2A^WT^ vs CAMK2A^LSL^;CAG-Cre^ESR^ p > 0.999, CAMK2A^LSL^ vs CAMK2A^LSL^;CAG-Cre^ESR^ p = 0.867). **C)** Paired-pulse facilitation, a form of presynaptic-dependent short-term plasticity (one-way ANOVA with Bonferroni’s post-hoc analysis calculated per interval. 10 ms: F_(3,107)_ = 0.16, p = 0.922; 25 ms: F_(3,107)_ = 5.34, p = 0.0018, CAMK2A^WT^ vs CAMK2A^WT^;CAG-Cre^ESR^ p > 0.999, CAMK2A^WT^ vs CAMK2A^LSL^ p > 0.999, CAMK2A^WT^ vs CAMK2A^LSL^;CAG-Cre^ESR^ p > 0.0686, CAMK2A^LSL^ vs CAMK2A^LSL^;CAG-Cre^ESR^ p = 0.0012; 50 ms: F_(3,107)_ = 2.88, p = 0.0393, CAMK2A^WT^ vs CAMK2A^WT^;CAG-Cre^ESR^ p > 0.999, CAMK2A^WT^ vs CAMK2A^LSL^ p > 0.999, CAMK2A^WT^ vs CAMK2A^LSL^;CAG-Cre^ESR^ p > 0.0572, CAMK2A^LSL^ vs CAMK2A^LSL^;CAG-Cre^ESR^ p = 0.1103; 100 ms: F_(3,107)_ = 2.35, p = 0.0763; 200 ms: F_(3,107)_ = 2.09, p = 0.1058; 400 ms: F_(3,107)_ = 1.18, p = 0.3210). **D)** Normalised fEPSP slope after induction of LTP with 100 Hz stimulation over time, insets show representative responses before (black) and after (coloured) LTP induction. **E)** Mean normalised fEPSP slope in the final 10 min of recording (one-way ANOVA with Bonferroni’s post-hoc analysis, F_(3,73)_ = 5.55, p = 0.0017, CAMK2A^WT^ vs CAMK2A^WT^;CAG-Cre^ESR^ p > 0.999, CAMK2A^WT^ vs CAMK2A^LSL^ p = 0.0090, CAMK2A^WT^ vs CAMK2A^LSL^;CAG-Cre^ESR^ p > 0.999, CAMK2A^LSL^ vs CAMK2A^LSL^;CAG-Cre^ESR^ p = 0.0221). Data represents mean ± SEM, n = 23-30 slices of 5-6 mice for each group. * p < 0.05, ** p < 0.01.

### *Adult* Camk2a *reinstatement rescues intrinsic behavioural phenotypes*

Most of the phenotypes tested above depend on hippocampal plasticity, which appears to not have a critical developmental window, but remains important throughout life (12, 16, 20, 31). Hence, we also wanted to assess the role of CAMK2A in more intrinsic behaviour, which has been shown to have an early critical developmental time window during which alterations can cause irreversible damage (16). For this purpose, we performed a behavioural battery of different tests, previously shown to be sensitive to a critical treatment window to obtain full reversal (16, 32–34). Similar to the previous experiments, gene reinstatement in this cohort was induced in adult 11-19-week-old mice and 5 weeks later a behavioural battery was performed. No behavioural phenotype was found in CAMK2A^LSL^ mice in the open field, rotarod, or marble burying test (data not shown). However, in both the nest building and forced swim test we found CAMK2A^LSL^ mice to show significantly altered behaviour compared to CAMK2A^WT^ mice. In the nest building paradigm, CAMK2A^LSL^ mice used less material to build their nest compared to the other genotypes and in the forced swim test, CAMK2A^LSL^ mice showed decreased immobility compared to CAMK2A^WT^ mice (Figure 4A and B). Surprisingly, both phenotypes were rescued in the CAMK2A^LSL^;CAG-Cre^ESR^ mice as they showed similar behaviour compared to CAMK2A^WT^ mice, suggesting that although absence of CAMK2A during early developmental periods affects these intrinsic behaviours, adult CAMK2A expression is beneficial to rescue these behaviours (Figure 4A and B). This could indicate that the impaired circuits that underlie the nest building and forced swim test deficits in *Camk2a* mice, are distinct from the circuits that underlie these deficits in *Ube3a* mice (16).

**Figure 4:**
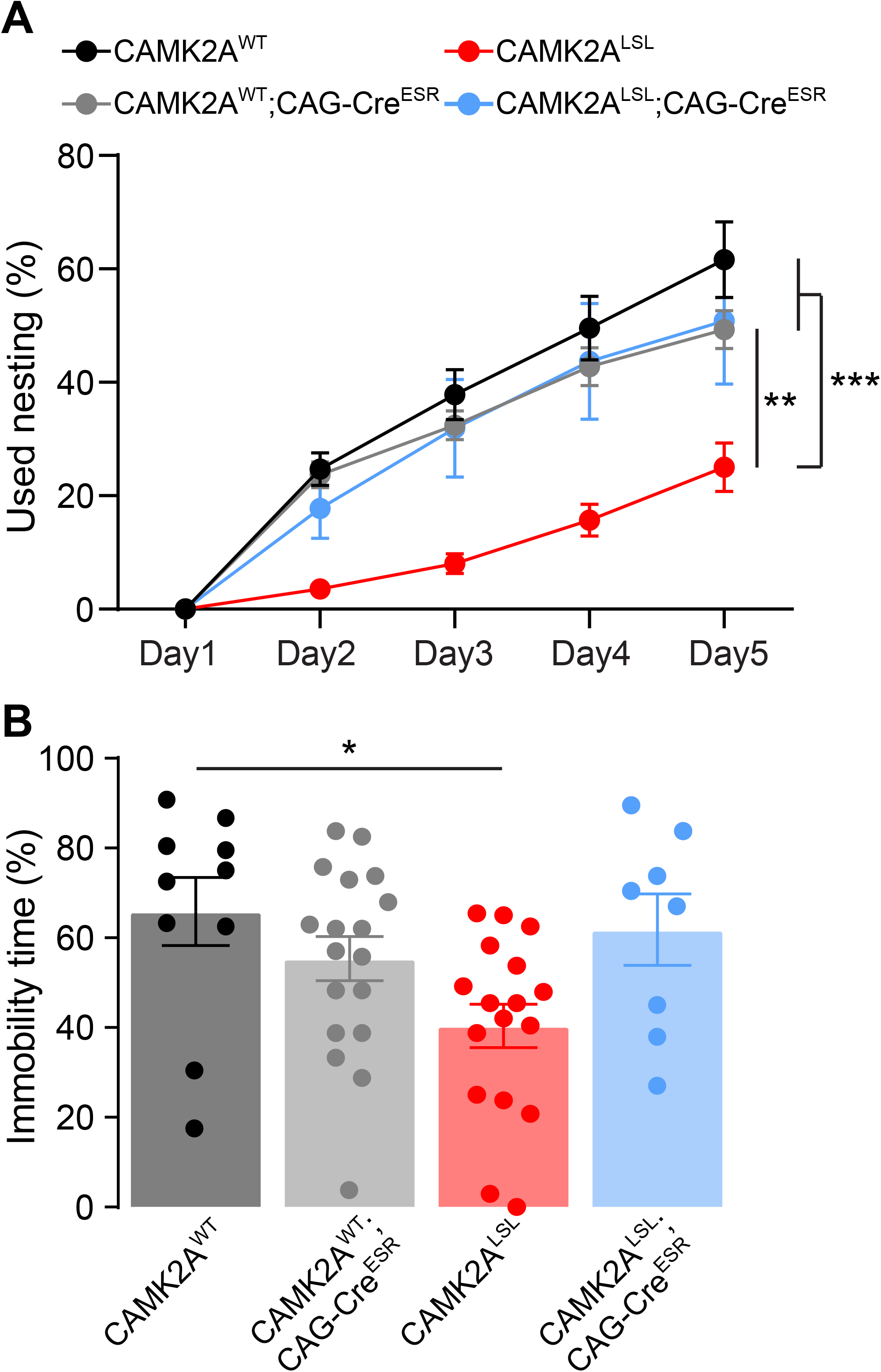
Adult CAMK2A expression rescues intrinsic behaviour in nest building and forced swim test. **A)** Amount of nest material used each day during nest building test over 5 days (two-way repeated measures ANOVA with Bonferroni’s post-hoc analysis on genotype. Interaction time x genotype: F_(12,196)_ =8.79, p < 0.0001; genotype: F_(3,49)_ =12.58, p < 0.0001, CAMK2A^WT^ vs CAMK2A^WT^;CAG-Cre^ESR^ p > 0.999, CAMK2A^WT^ vs CAMK2A^LSL^ p < 0.0001, CAMK2A^WT^ vs CAMK2A^LSL^;CAG-Cre^ESR^ p > 0.999, CAMK2A^LSL^ vs CAMK2A^LSL^;CAG-Cre^ESR^ p = 0.0020. **B)** Amount of time mice spent immobile in forced swim test (one-way ANOVA with Bonferroni’s post-hoc analysis, F_(3,49)_ = 3.68, p = 0.0182, CAMK2A^WT^ vs CAMK2A^WT^;CAG-Cre^ESR^ p > 0.999, CAMK2A^WT^ vs CAMK2A^LSL^ p = 0.0267, CAMK2A^WT^ vs CAMK2A^LSL^;CAG-Cre^ESR^ p > 0.999, CAMK2A^WT^;CAG-Cre^ESR^ vs CAMK2A^LSL^ p = 0.2649, CAMK2A^LSL^ vs CAMK2A^LSL^;CAG-Cre^ESR^ p = 0.1233). Data represents mean ± SEM, n = 8-18 mice per group. * p < 0.05, ** p < 0.01, *** p < 0.001.

## Discussion

With the current advance in genetic diagnosis, causing a quickly expanding list of rare neurodevelopmental disorders and the upcoming possibility to treat these disorders with different forms of gene therapy, such as antisense oligonucleotides, the need to understand the optimal timing for treatment becomes increasingly important. CAMK2-related disorder is one of these recently discovered disorders, and our aim in this study was to understand whether a developmental critical time window exists for CAMK2A to be expressed for normal brain development and function. For this purpose, we generated an inducible CAMK2A knockout model, where we could restore CAMK2A expression in adulthood and assessed the behaviour. Together, our results imply that loss of CAMK2A during development does not cause irretrievable distortion to neural circuits in the brain, as we fully rescue the behavioural and electrophysiological phenotypes assessed here.

As far as we know this is the only NDD mouse model described in literature, where all phenotypes can be fully normalised upon adult gene reinstatement. In a subset of the NDD mouse models, adult gene reinstatement rescues some but not all behavioural deficits (e.g. upon reinstatement of SHANK3 (15)), rescues observed phenotypes partially (e.g. upon reinstatement of MECP2 (19)), or only rescues the behavioural deficits if reinstated at a juvenile age (e.g. upon reinstatement of UBE3A or SYNGAP1 (16, 18)). The reverse is also true, as only adult deletion of *Camk2a* completely phenocopies the deficits observed in germline knockout mice (24), whereas adult deletion in other NDD models do either not or only partially copy the observed behavioural abnormalities (e.g. upon *Syngap1, Mecp2* or *Ube3a* deletion (31, 33, 35)), or only show the phenotypes upon juvenile deletion (e.g. upon *Ube3a* deletion (33)), proving a critical window for protein expression of this gene.

CAMK2A is a key player in NMDA-receptor dependent long-term potentiation. Recently, the group of Amy Ramsey published the effect of adult reinstatement of GRIN1 on behaviour, another NDD mouse model in the same pathway as CAMK2A (14). Adult reinstatement of GRIN1 resulted in significant improvement in all behaviours, but not all were fully normalised. Social behaviour was fully rescued, as could be expected based on the critical window. However, cognitive behaviour, tested with a puzzle box, was not fully rescued, nor were the more intrinsic novelty induced locomotion or anxiety-like behaviour. Surprisingly, our mouse model does show full recovery for anxiety-like and intrinsic behaviour. These combined results indicate that even for genes from the same pathway (e.g. NMDA-receptor dependent plasticity), the effect of reinstatement in adult age can be different, further emphasising the importance of testing this for every gene found to cause an NDD. One of the reasons for this difference could be the temporal expression. Whereas CAMK2A starts to be expressed at postnatal day 1, GRIN1 is already expressed at early time points (from E9.5) of prenatal development (36).

The results presented here suggests that CAMK2A does not play a role during development. However, it was recently shown that deletion of *Camk2a* and *Camk2b* simultaneously from germline, is lethal at P1, the moment that CAMK2A starts to be expressed (37). With the *Camk2a* and *Camk2b* single knockouts being completely viable, this finding does suggest an important early postnatal developmental role also for CAMK2A, but one that can be compensated for by CAMK2B. Our results now imply that this compensation is of crucial importance for early postnatal brain development, preventing irreversible alterations in the neuronal circuit and therefore allowing for full phenotypic recovery in the adult mice.

To conclude, we show that CAMK2A expression is required during learning or other types of behaviour, independent from its expression throughout development. This could be of crucial importance for patients that carry *CAMK2A* loss-of-function mutations (7, 8), and opens doors towards potential use of therapies that increase CAMK2A expression.

## Methods

### Generation of mouse line

To generate the CAMK2A^LSL^ mouse model, the construct used to generate the *Camk2a* floxed mouse line was used as a starting point (24). From this construct the 5’ flanking arm of exon 2 and the 3’ flanking arm containing exon 2 of *Camk2a* were obtained and cloned in a vector containing the cassette of *Camk2a*-exon2 fused to tdTomato, followed by the woodchuck hepatitis virus post-transcriptional regulatory element (WPRE), flanked by LoxP sites, and a neomycin cassette for positive selection, flanked by Frt sites. All exonic sequences were sequenced to verify that no mutations were introduced accidentally. The targeting construct was linearized and electroporated into E14 ES cells (derived from 129P2 mice). Cells were cultured in BRL cell-conditioned medium in the presence of leukemia inhibitory factor. After selection with G418 (200 *μ*g/ml), targeted clones were identified by PCR (long-range PCR from neomycin resistance gene to the region flanking the targeted sequence). A clone with normal karyotype was injected into blastocysts of C57BL/6J mice. Male chimeras were crossed with female C57BL/6J mice and resulting offspring was further backcrossed into the C57BL/6J background. To obtain the mice used for experiments, heterozygous CAMK2^LSL^ females (15 times backcrossed in C57BL/6J) were crossed with Tg(CAG-cre/Esr1*)5Amc (CAG-Cre^ESR^) male mice (MGI:2182767)(17 times backcrossed in 129S2/SvPasCrl). Of the F1 offspring, the heterozygous CAMK2A^LSL^;CAG-cre^ESR^ were crossed with heterozygous CAMK2A^LSL^ mice, obtaining the resulting experimental group: CAMK2A^LSL^, CAMK2A^LSL^;CAG-Cre^ESR^, CAMK2A^WT^ and CAMK2A^WT^;CAG-Cre^ESR^. Mice of both sexes were used for the experiments. During the experiments, the experimenter was blind for the genotypes. All mice were group housed at 22±2 °C, except for the nest building test when they were single caged. They were on a 12/12h light/dark cycle with food and water available *ad libitum*. All experiments were performed during the light phase. All animal experiments were conducted in accordance with the European Commission Council Directive 2010/63/EU (CCD project license AVD101002017893), and all described experiments and protocols were subjected to ethical review (and approved) by an independent review board (IRB) of the Erasmus MC.

### Tamoxifen treatment

To induce Cre-mediated deletion, mice received intraperitoneal injections of 0.1 mg Tamoxifen per gram body weight for four consecutive days at 11-19 weeks old. Tamoxifen was freshly dissolved at a concentration of 20 mg/ml in sunflower oil beforehand.

### Immunohistochemistry

Adult mice were transcardially perfused with paraformaldehyde (4%) to fixate tissue and brains were extracted. Gelatin-embedded slices of 40 µm thick were blocked in 3% H_2_O_2_ in PBS, rinsed, and for antigen retrieval incubated in sodium citrate (10 mM) at 80 °C for 20 min. The slices were again rinsed and placed in blocking solution (10% normal horse serum and 0.5% Triton X-100 in PBS) for 1h at room temperature and incubated overnight at 4 °C in blocking solution with AffiniPure Fab Fragment (donkey anti mouse, 1:200, catalog #rid_000053, Jackson ImmunoResearch). The next day, slices were incubated for 48-72h with primary antibodies (anti-CAMKII alpha (6G9 clone), 1:100, NB100-1983, Novus Biologics; anti-RFP, 1:5000, 600-401-379, Rockland) shaking delicately at 4 °C. Secondary antibodies (Alexa, 1:200, Jackson ImmunoResearch) were incubated for 2h at room temperature. Finally, slices were incubated with 40,6-diamidino-2-phenylindole solution (DAPI, 1:10,000, Invitrogen) for 10 min and mounted in Mowiol medium. Images were acquired with a confocal microscope (LSM700, Zeiss).

### Western blot

Mice were anaesthetized with isoflurane. Cortical and hippocampal tissue was isolated and snap frozen in liquid nitrogen. Samples were sonicated in lysis buffer (0.1 M Tris-HCl [pH 6.8], 4% SDS), containing protease inhibitor cocktail (P8340, Sigma). Protein concentration of the supernatant was determined using the BCA protein assay (Pierce™) and 15 µg was loaded on the gel (3450124, Bio-rad), which was next turbo transferred on nitrocellulose membrane (1704159, Bio-rad). Blots were probed with primary antibodies (anti-CAMKII alpha (6G9 clone), 1:20,000, NB100-1983, Novus Biologics; anti-RFP, 1:2000, 600-401-379, Rockland) overnight at 4 °C. Next day, blots were incubated with IRDye® 800CW Goat anti-Mouse and 680LT Goat anti-Rabbit secondary antibodies (Licor, 1:15,000) for 1 h at room temperature. Blot were imaged using an Odyssey® Imager (Licor) and analysed using Image Studio™ Lite (Licor).

### Immunoprecipitation

Cortical tissue was homogenised in lysis buffer (1mM NaHCO_3_, 1 mM MgCl_2_, 0.32 M sucrose, 10 mM Hepes [pH 7.4]) containing protease inhibitor cocktail (P8340, Sigma). Supernatant, containing soluble proteins, was pre-cleared with Protein A agarose beads (P3476, Sigma). An ‘input’ sample was set aside and 900 µl of supernatant was incubated overnight with beads cross-linked to CAMK2A primary antibody (10 µg, NB100-1983, Novus Biologics) rotating end-over-end at 4 °C. After washing (150 mM NaCl, 0.1% Triton X-100, 25 mM Hepes [pH 7.4]), protein was eluted by boiling for 5 min at 95 °C. Input, unbound and bound IP samples were handled by western blot, which was probed for CAMK2A (1:20,000, NB100-1983, Novus Biologics), CAMK2B (1:10,000, 13-9800, Invitrogen) and Actin (1:20,000, MAB1501R, Millipore).

### Behaviour

All experiments were conducted during the light phase of the light/dark cycle and experimenters were blind to genotype. Mice were handled beforehand.

#### Morris water maze

Mice were trained to find a submerged platform (1 cm submerged, 11 cm in diameter) in a circular pool (1.2 meter in diameter) for 5 days. Trials consisted of placing the mouse 30 sec on the platform and releasing it at a pseudorandom location in the water maze. If the mouse did not locate the platform within 60 sec, the experimenter gently guided the mouse to it. The same procedure was then repeated a second time. Latency to find the platform was manually scored. A probe trial was carried out on day 6, in which the platform was removed after the mouse was initially placed there for 30 sec. During the probe trial the mouse was tracked using EthoVision® software (Noldus®). Throughout the experiment the water was made opaque with white paint and kept at a temperature of 25-26 °C.

#### Fear conditioning

Mice were placed in a soundproof box (26 × 22 × 18 cm; San Diego Instruments) with a grid floor, white light turned on and a camera monitoring the behavioural activity. The training session consisted of 200 sec, wherein after 150 sec a conditioned stimulus of a tone (20 sec, 85 dB) was presented followed by the unconditioned stimulus of a foot shock via the grid floor (1.0 mA, 2 sec) after which the mice were left in the box for the remaining 28 sec. 24h later, contextual memory was monitored for 180 sec by placing the mice in the same box. Another 24h later, cued memory was monitored for 220 sec by placing the mice in an adapted version of the box and after 120 sec presenting them with the conditioned stimulus for 100 sec. The box was adapted by removing the grid floor, placing plexiglass walls in a triangular shape, turning the light off, and presence of an acetonic odour. Freezing behaviour was analysed with Video Freeze® software (Med Associates Inc) and started when the mouse showed no activity for 1.00 sec.

#### Nest building

The mice were single caged 5-7 days before the start of the experiment. Used nest building material was removed from the cages and replaced by 11 g compressed extra-thick blot filter paper (Bio-rad). Every 24h for 5 days, unused material was weighted.

#### Forced swim test

Mice were placed in a glass cylinder (diameter of 18 cm, height of 27 cm) filled with water up to 15 cm (26±1 °C) for 6 min and recorded with a camera. After 2 minutes of habituation, the mice were scored manually for immobility for 4 minutes. Immobility was defined as no activity other than that needed for the mouse to remain afloat and keep its balance.

### Electrophysiology

Mice were decapitated under anaesthesia of isoflurane and brains were rapidly removed. Sagittal slices of 400 µm were cut with a vibratome (PELCO easiSlicer™, Ted Pella) in ice cold oxygenated (95%) and carbonated (5%) artificial cerebral spinal fluid (ACSF; in mM: 120 NaCl, 3.5 KCl, 2.5 CaCl2, 1.3 MgSO4, 1.25 NaH2PO4, 26 NaHCO3, and 10 D-glucose). Hippocampi were isolated and allowed to recover in oxygenated and carbonated ACSF at room temperature for 90 min. Slices were placed in a submerged chamber with an ACSF flow of 2 ml/min at 30 °C degrees. Bipolar platinum/iridium electrodes (Frederick Haer) were placed in the Shaffer collateral CA1-CA3 region for stimulation and in the dendrites of CA1 pyramidal cells for recording. Slices were left to habituate to the electrode placements for 30 min, before the onset of the experiment. Stimulations were at 1/3^rd^ max fEPSP strength and lasted 100 µs. Paired-pulse facilitation was measured by three rounds of two sequential stimulations at intervals of 10 – 25 – 50 – 100 – 200 – 400 µs. For the LTP experiment, responses were recorded at 1 Hz. After 10 min baseline, LTP was induced by tetanic stimulation of 100 Hz for 1 sec and recorded for 60 min. During analysis, responses were normalised to the baseline. Recordings that showed an unstable baseline were excluded.

### Statistics

Statistical significance was determined using one-way or two-way ANOVA test with Bonferonni post hoc comparison in PRISM software (Graphpad). P-values < 0.05 were considered significant. Exact p-values are indicated in the figure legends, CAMK2A^WT^;CAG-Cre^ESR^ vs experimental groups are only represented if the significance is not equal to CAMK2A^WT^. When CAMK2A^LSL^ showed a phenotype compared to CAMK2A^WT^ mice, significance between all groups is indicated in the figures.

### Study approval

All animal experiments were conducted in accordance with the European Commission Council Directive 2010/63/EU (CCD project license AVD101002017893), and all described experiments and protocols were subjected to ethical review (and approved) by an independent review board (IRB) of the Erasmus MC

## Author contributions

GW and YE designed the experiments. LP generated the mouse line. MAJ conducted and analysed the electrophysiological experiments, all other experiments were conducted and analysed by GW, JK, IW and PR. PR and GW wrote the manuscript.

## Acknowledgements

We would like to thank Armando Schoonbrood for technical assistance.

Address correspondence to: Ype Elgersma, Department of Clinical Genetics, Tel: +31107043337; Or to Geeske van Woerden, Department of Clinical Genetics and Department of Neuroscience, Tel: +31107032695, Erasmus Medical Center, Rotterdam, Dr. Molewaterplein 40, 3015 GD, The Netherlands.

